# Thiel Embalming Pre-Frozen Cadavers: A Mouse Model

**DOI:** 10.1101/523514

**Authors:** Murad AlShehry, Mostafa Kandil, Raed Alzahrani, Roger Soames

## Abstract

Thiel embalming has been gaining a great deal of worldwide attention. Its long-lasting life-like quality enables multiple applications in surgical training programs. The standard process of Thiel embalming, which involves perfusion of embalming fluid in fresh cadavers through cannulation of the arterial and venous systems could be difficult in some areas of the world where fresh body donation rates are low or banned. This study will assess the ability to Thiel embalm fresh frozen mice as an animal model as a solution for the lack of fresh cadavers. The ability of Thiel embalming frozen tissue would enable areas in the world where they do not have access to recently diseased donated cadavers. The study was ethically approved King Fahad Medical City Institutional Review Board. The 20 Mice cadavers were ethically gained from the university of Dundee animal house. They were euthanized ethically by carbon dioxide. They were handled by following the university’s ethical code. The mice were split into 4 groups with 5 mice for each group, where group 1 is fresh unfixed. Group 2 standard Theil embalmed. Group 3 defrosted one-week frozen mice then standard embalmed. group 4 perfused with Theil fluid, frozen, defrosted then set in the embalming tank. The embalming validation tests were made by visual gross and histological changes in the gastric, renal and muscular tissue. These tissues were chosen to test the penetration of the embalming solution. The results show similarity in all groups with difference of internal gross changes in the pre-frozen cadavers. Histological changes were similar in all embalmed groups meaning that the embalming method has been successful from a histological point of view.

## Introduction

Thiel embalming has been gaining increasing worldwide attention, not least because of its long-lasting life-like quality enabling multiple applications in surgical training programs. The process of Thiel embalming involves perfusion of embalming fluid in fresh cadavers through cannulation of the arterial and venous systems; however this could be difficult in some areas of the world where fresh body donation rates are low or banned. The current study assesses the ability to Thiel embalm fresh frozen cadavers, using a mouse model, as a solution to the lack of fresh cadavers.

Wolff^1^ reported that Thiel embalming is a unique technique ideally suited to teach surgical flap raising and microvascular suturing on human material. They commented that the technique permitted flap raising procedures to be undertaken under realistic conditions, i.e. similar to those in the living. They also reported that vessels and nerves up to a diameter of 1 mm could be exposed and dissected enabling microvascular suturing as in fresh specimens even after several weeks. Dissection of soft and hard tissues, as well as performing implantological procedures, was reported by Hölzle ^2^ to be very life-like in Thiel embalmed cadavers, noting that the maxillary sinus membrane, mucosa, bone and nerves could be exposed, dissected, drilled and sutured several weeks after embalming, being similar to fresh specimens and the living. These authors emphasise that Thiel embalming is ideally suited to teaching oral surgery and implantology on human material.^2^ Odobescu ^3^ assessed the utility of a Thiel arterial model in microsurgical research by comparing interrupted horizontal mattress (HM) sutures to simple interrupted (SI) sutures in human vessels. They evaluated vessel patency, leak and stricture using angiography, and vessel wall architecture using light and scanning electron microscopy (SEM): technique speed was also assessed. The authors concluded that the artery research model was promising, with the HM microvascular suture requiring further *in vivo* validation.^3^

Following hands-on training courses on Thiel embalmed cadavers (THC) in colon, bariatric and vascular surgery Giger ^4^ stated that training on such cadavers could be an excellent additional model to teach advanced bariatric, hernia and colon surgery. However, they noted that it is important to define which training model (THC, anesthetized animals, virtual computer training, etc.) was the most appropriate for the skill or procedure being taught. Hassan ^5^ later compared Thiel embalmed cadavers with formalin embalmed cadavers and porcine models in the surgical repair of flexor tendons, with each model being compared on a 5-point scale to assess tissue quality, surgical approach and identification of structures. Thiel cadavers rated consistently higher compared with the formalin and porcine models and demonstrated the benefit of Thiel embalming retaining tissue flexibility allowing testing the repaired flexor tendon in a realistic anatomical model. In the same year Willaert ^6^ reported on the development of a circulation model in Thiel embalmed pig kidneys and raised a number of issues, among which were the ethics of pursuing methods to preserve animal cadavers for use in surgical techniques in an attempt to reduce the use of living animals for teaching. They reported that the quality of tissue preservation in Thiel embalmed cadavers varied, but that arterial embalming of kidneys resulted in successful preservation due to complete parenchymatous spreading. Nevertheless, they concluded that more research is required to determine whether other factors affect embalming quality.^6^

The ability to Thiel embalm frozen tissue could enable areas in the world where there is no access to donated cadavers to develop and run surgical training courses and programs.

## Methods

Laboratory mice from a single strain were used in the current study: mice were selected on the basis of their size, allowing faster profusion of the active ingredients of the Thiel solution into the whole animal. The mice were supplied by the College of Life Sciences University of Dundee, euthanized ethically by carbon dioxide, and handled following guidelines in the university’s ethical code.

The 20 mice used in the study were divided into 4 groups of 5 mice each.

- **Group 1**: control group of fresh unfixed mice used as a reference for gross and histological assessment.
- **Group 2**: standard Thiel embalmed mice used as a Thiel control.
- **Group 3**: perfused then frozen at −20^0^ C in bags with fluid for 1 week then Thiel embalmed.
- **Group 4**: pre-frozen at −20^0^ C for 1 week then thawed for 4 hours at room temperature prior to undergoing Thiel embalming.

Groups 2, 3 and 4 were labeled with colored cable ties attached to the base of the tail to distinguish between groups.

The Thiel embalming protocol used in this study was advised by the Centre of Anatomy and Human identification at the University of Dundee, with the protocol employed being adopted from the studies of Zeller ^7^ and Gage ^8^. Standard arterial perfusion through either the femoral or common carotid arteries cannot be applied in mice, therefore injection into the left ventricle was employed.^(7,8)^ By exposing the thorax the left ventricle was accessed with minimal dissection. Each animal to be perfused was secured to a cork board to prevent inadvertent movement. A skin incision was made between the top of the abdomen and neck, following which two horizontal incisions from the thorax to the axilla were made: the ribcage was sectioned so that its anterior aspect hinged like a flap. An intravenous bag filled with the Thiel arterial infusion solution, prepared by the Centre for Anatomy and Human identification, was placed at a higher level than the animal to facilitate gravity perfusion: the composition of the arterial perfusion fluid is given in Table 1. The IV bag was connected to a size 18 IV needle that had a roller clamp to control the flow of the solution: the needle was introduced into the left ventricle, following which the roller clamp was slowly released to gradually increase the flow pressure of the perfusion fluid. The amount of fluid needed for complete perfusion was between 15 and 20mL depending on the size of the animal.

**TABLE 1.** Composition of 1.3 litres arterial infusion Thiel embalming solution.^9^

| Materials | Amount |
| --- | --- |
| Hot tap water | 680 mL |
| Boric acid | 20g |
| Ammonium nitrate | 168g |
| Potassium nitrate | 42g |
| Sodium sulphite | 70g |
| Propylene glycol | 250 mL |
| 1:10 chlorocresol in glycol | 50 mL |
| Formaldehyde solution 8.9% | 210 mL |
| Morpholine | 15 mL |
| Ethanol | 100 mL |
| Total Volume | 1.3 L |

This method enabled the rapid delivery of embalming solution to the tissues via the circulation. Tail and limb tremors, whole body swelling, loss of a dark color of the liver and arterial solution leaking from the mouth and nose were taken as signs of complete perfusion. Following perfusion each animal in groups 2, 3 and 4 was submerged in 4 L of Theil arterial infusion solution for 5 days to facilitate full tissue fixation.

On completion of the embalming process mice in all three groups were dissected to determine the extent of fixation, as well as the appearance and flexibility of the Thiel embalmed specimens as reported by Eisma.^9^

One sample of skeletal muscle tissue, approximately 0.5 cm^3^, from the left hamstring, as well as tissue samples from the kidney and fundus of the stomach, were harvested from each animal. These tissues were selected to represent different depths within the body and act as an indication of the level of Thiel perfusion within the animal. The samples were processed using a standard paraffin embedding protocol, i.e. fixation with formalin, ethanol dehydration, clearing with xylene and finally embedding in hot paraffin wax: only Group 1 underwent fixation with formalin as Groups 2, 3 and 4 were already fixed using Thiel. The resulting tissue blocks were sectioned at 7μm, mounted on slides and stained with Hematoxylin and Eosin (H&E). The slides were analyzed and comparisons made between the groups as well as with the literature.

Embalming validation was assessed by visual gross and histological changes observed in the gastric, renal and muscular tissue.

The study received ethical approval from the King Fahad Medical City Institutional Review Board.

## Results

### Perfusion

Perfusion in groups 2 and 3 resulted in the animals swelling followed by clear embalming solution leaking from the nose and mouth. The liver and lungs became paler showing that the embalming fluid had perfused these organs. In group 4, however when perfusion was complete the leaking fluid was mixed with blood: the liver and lungs in this group did not change colour to the same extent as in the fresh cadavers.

### Gross Analysis

Following 5 days submersion the initial gross sign of complete embalming was the loss of fur: any remaining fur was removed by rubbing and then rinsing in a still water bath. Following fur removal, the mice were returned to the arterial infusion fluid for a further day to recover the lost embalming fluid before storing in plastic bags.

Group 2 and 3 showed similar effects of embalming, both at a gross level and histologically. However, in group 4 the gastrointestinal system appeared to have deteriorated to a much greater extent than in groups 2 and 3 (Figure 1): this was observed as the intestines being somewhat brittle. This gastrointestinal system change is probably the result of slow bioactivity of the gut flora and enzymes while the animals were frozen. In addition, group 4 also showed a greater degree of tissue deterioration at the gross level only.

**Figure 1:**
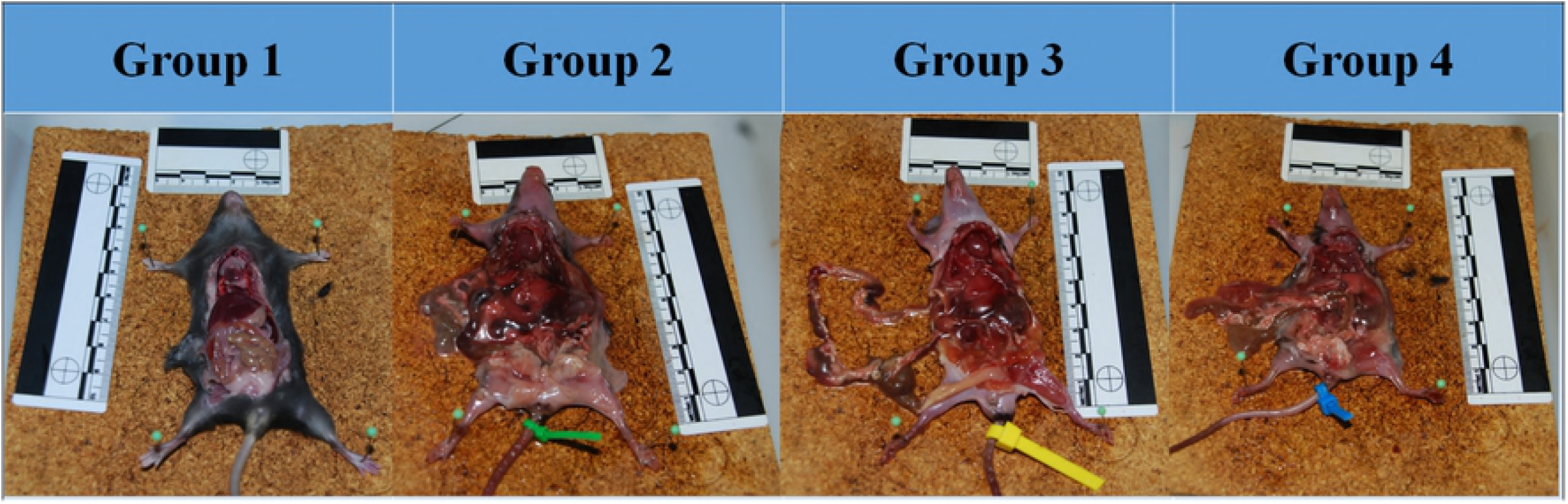
Gross dissection of the 4 groups, from left to right Group 1, fresh; Group 2, standard Thiel embalmed; Group 3, perfused, frozen then submerged in Thiel fluid; and Group 4, frozen then thawed followed by standard Thiel embalming.

### Histological Analysis

The histological sections showed complete embalming for all animals in groups 2, 3 and 4, observed microscopically by the absence of hematoxylin stained cell nuclei compared with group 1 (non-embalmed tissue). The cell nuclei were stained dark purple by the hematoxylin and surrounded by pink endoplasm, stained by the eosin. The majority of the Thiel embalmed groups showed no signs of purple staining indicating completely embalmed tissue (Figure 2).

**Figure 2:**
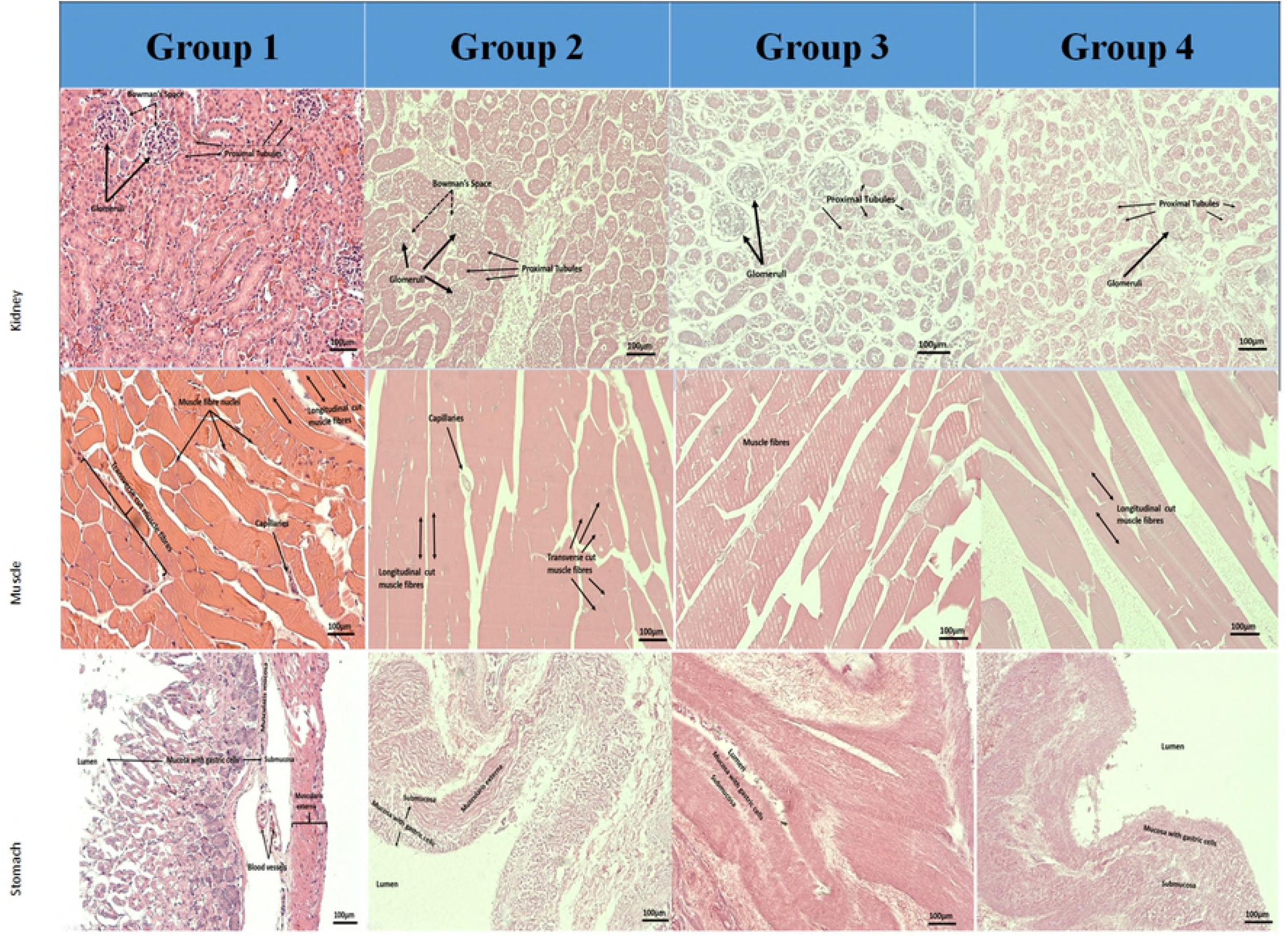
H&E staining of Kidney, muscle and stomach tissues for Group 1 (fresh unfixed) Group 2 (Thiel embalmed) Group 3 (perfused then frozen at −200 C) and Group 4 (pre-frozen at −200 C) at magnification of ×10.

## Discussion

### Arterial perfusion technique

In the animal model used the initial dissection was minimal to preserve other tissues and observe the flow of embalming fluid through the circulation. It was noted that to achieve a good flow rate relatively little pressure was required; however care has to be taken to maintain an adequate pressure to keep the needle in place at the base of the heart. Too much fluid pressure tends to push the needle out of the left ventricle.

### Gross Findings

Thiel embalming involves three main processes: fixation, disinfection and preservation. Applying the three processes with specific solutions results in the preservation of organs and tissues with their natural color.^(10,11)^ The solutions used in Thiel embalming give muscles a more reddish colour^12^: it is the absorption of nitrate in the formation of nitrosomyoglobin which gives muscles a red colour.^12^ According to Hayashi ^13^ Thiel embalmed cadavers maintain joint and muscle flexibility and mobility.

Reports in the literature and the author’s experiences of Thiel embalmed cadavers confirms the successful embalming in groups 2, 3 and 4. The cadaver was flexible demonstrating the Thiel flexibility properties, the colour of the muscles and fat tissue were similar to the texture of the tissues of Thiel embalmed cadavers. Furthermore, in spite of being stored in sealed bags with little embalming fluid for up to 2 months there were no visible signs of decay, loss of flexibility or colour. The internal organs, such as the spleen, heart, kidney and liver, had similar consistency and texture to those in human Thiel embalmed cadavers at the Centre for Anatomy and Human Identification University of Dundee.^9^

The gastrointestinal tract was intact and no further decay was observed in groups 2 and 3; however group 4 showed a brittle gastrointestinal tract. Willaert ^6^ reported that animal renal tissue had a range of qualities following embalming. It is postulated that gastrointestinal tissue could also be variable in quality, thus the observations in group 4. It is also possible that given the small size of the gastrointestinal tract in mice that the tissue showed early signs of deterioration. In larger animals, including humans, the gastrointestinal tract could remain intact even after freezing for periods longer than 1 week.

### Histological findings

Histological examination of sections from groups 2, 3 and 4 revealed no evidence of cell nuclei, being similar to the observations of Tennent ^14^. Fragmentation of muscle proteins was observed in groups 2, 3 and 4, similar to the observations of Benkhadra.^15^ These observations indicate that the tissue had been modified by the embalming process and was therefore successfully embalmed at a histological level. It was noted that the group 1 samples were well stained showing the cellular structures in great detail. Unlike group 1, groups 2, 3 and 4 exhibited paler sections suggesting that the histological stains did not bind as well as in group 1. This is most likely due to the increased acidity resulting from the large amounts of boric acid in the submersion solution.

### Limitation

There are current regulations to the use and disposal of ammonium nitrate, with increased governmental restrictions in many countries due to its potential use in terrorist attacks: improper storage could also lead to damage to lives and property.^16^

Excessive boric acid concentration in the arterial perfusion fluid is assumed to be responsible for the failure to stain adequately some histological sections. It is recommended that lower concentrations of boric acid for embalming small animals such as mice.

### Strength

The rationale behind this study was the development of a novel method that could be employed to Thiel embalm fresh frozen cadavers. The study has shown that small blood vessels can accommodate the introduction of embalming fluid after a period of 1 week following death.

### Future work

The current pilot study on embalming fresh frozen cadavers has shown that it would be feasible to Thiel embalmed fresh frozen cadavers in Saudi Arabia, which has a huge demand for wet-laboratory surgical training courses/programs. The availability of flexible cadavers would increase the extent of professional training and thereby improve surgical skills. Ultimately this would improve the quality of medical care in the region.

## Conclusion

This study has successfully shown that Thiel embalming works well on frozen mice cadavers. Although the gross quality of the viscera appeared to be less robust in group 4, which had a poorer quality than in groups 2 and 3; nevertheless, this will enable fresh frozen cadavers to be Thiel embalmed opening the way for future cadaveric surgical training courses. The success of Thiel embalming of fresh frozen cadavers will help alleviate the lack of access to cadavers in countries where body donation is prohibited.

Thiel embalming enables cadaver preservation to facilitate medical and surgical teaching on the human body as a whole. In countries where fresh body donations are prohibited, such as Saudi Arabia, access to Thiel embalmed cadavers would be difficult.^17^ To overcome this challenge two scenarios are proposed. The first, based on the observations in group 3, where cadavers are perfused at source and then transported to the destination country where fresh cadavers are prohibited. After clearance by local authorities, the cadaver can then be submersed in Thiel tank fluid for 2months to complete the embalming process.^9^ The second scenario is based on the observations of group 4, where cadavers are fresh frozen in the source country, transported in temperature-controlled containers, thawed and embalmed at the receiving institute. Even if the time taken to transport the cadaver is weeks rather than days this should not influence the subsequent Thiel embalming process.

## Acknowledgement

The authors would like to give thanks the Research Center in King Fahad Medical City for funding this study through the Intramural fund #IRF 017-034. Additional thanks to the Centre for Anatomy and Human Identification at the University of Dundee for their support, use of facilities use and technical expertise, Professor Tracey Wilkinson, Amanda Hunter, Samantha Skene and Sheryl Paton.

## References

1. Wolff K, Kesting M, Mücke T, Rau A, Hölzle F. Thiel embalming technique: A valuable method for microvascular exercise and teaching of flap raising. Microsurgery. 2008;28(4):273–278.

2. Hölzle F, Franz E, Lehmbrock J, Weihe S, Teistra C, Deppe H et al. Thiel Embalming Technique: A Valuable Method for Teaching Oral Surgery and Implantology. Clinical Implant Dentistry and Related Research. 2009;14(1):121–126.

3. Odobescu A, Moubayed S, Harris P, Bou-Merhi J, Daniels E, Danino M. A new microsurgical research model using Thiel-embalmed arteries and comparison of two suture techniques. Journal of Plastic, Reconstructive & Aesthetic Surgery. 2014;67(3):389–395.

4. Giger U, Frésard I, Häfliger A, Bergmann M, Krähenbühl L. Laparoscopic training on Thiel human cadavers: A model to teach advanced laparoscopic procedures. Surgical Endoscopy. 2007;22(4):901–906.

5. Hassan S, Eisma R, Malhas A, Soames R, Harry L. Surgical simulation flexor tendon repair using Thiel cadavers: a comparison with formalin embalmed cadavers and porcine models. Journal of Hand Surgery (European Volume). 2014;40(3):246–249.

6. Willaert W, De Vos M, Van Hoof T, Delrue L, Pattyn P, D’Herde K. Understanding Thiel Embalming in Pig Kidneys to Develop a New Circulation Model. PLOS ONE. 2015;10(3):e0120114.

7. Zeller R. Fixation, embedding, and sectioning of tissues, embryos, and single cells. Current protocols in molecular biology. 2001;14.

8. Gage G, Kipke D, Shain W. Whole Animal Perfusion Fixation for Rodents. Journal of Visualized Experiments. 2012;(65).

9. Eisma R, Lamb C, Soames R. From formalin to thiel embalming: What changes? One anatomy department’s experiences. Clinical Anatomy. 2013;26(5):564–571.

10. Thiel W. The preservation of the whole corpse with natural color. Annals of anatomy. 1992 Jun;174(3):185–95.

11. Groscurth P, Eggli P, Kapfhammer J, Rager G, Hornung J, Fasel J. Gross anatomy in the surgical curriculum in Switzerland: Improved cadaver preservation, anatomical models, and course development. The Anatomical Record. 2001;265(6):254–256.

12. Janczyk P, Weigner J, Luebke-Becker A, Kaessmeyer S, Plendl J. Nitrite pickling salt as an alternative to formaldehyde for embalming in veterinary anatomy—A study based on histo-and microbiological analyses. Annals of Anatomy - Anatomischer Anzeiger. 2011;193(1):71–75.

13. Hayashi S, Homma H, Naito M, Oda J, Nishiyama T, Kawamoto A et al. Saturated Salt Solution Method. Medicine. 2014;93(27):e196.

14. Tennent S, Soames R, Felts P. Understanding and improving Thiel. Journal of Anatomy. 2012 Jul 1;221(1):85–6.

15. Benkhadra M, Bouchot A, Gérard J, Genelot D, Trouilloud P, Martin L et al. Flexibility of Thiel’s embalmed cadavers: the explanation is probably in the muscles. Surgical and Radiologic Anatomy. 2010;33(4):365–368.

16. Johnston L. Fertilizer Used In Terror Bombs [Internet]. Cbsnews.com. 2018 [cited 22 October 2018]. Available from: https://www.cbsnews.com/news/fertilizer-used-in-terror-bombs.

17. Habbal O. The state of human anatomy teaching in the medical schools of Gulf Cooperation Council countries: Present and future perspectives. Sultan Qaboos University Medical Journal. 2009 Apr;9(1):24.

